# A gapless telomere-to-telomere reference genome of *Ostreococcus tauri* RCC4221 with expanded annotation of medium-sized ncRNAs

**DOI:** 10.64898/2026.07.10.737489

**Authors:** Gaokang Liu, Laurie Bousquet, Helene Mayeur, Eric Manirakiza, Vladimir Daric, David Lopez-Escardo, Céline Noirot, Christophe Klopp, Sheree Yau, Marc Krasovec, Nigel Grimsley, Manuel Echeverria, Gwenaël Piganeau

**Author notes:** Corresponding author Gwenaël Piganeau, Sorbonne Université, university of Perpignan via domitia, CNRS, UMR 8176 LBBM, Observatoire Océanologique de Banyuls, avenue Pierre Fabre, Banyuls-sur-Mer, France. These authors contributed equally to this work (Gaokang Liu, Laurie Bousquet, Helene Mayeur).

## Abstract

Marine photosynthetic microbes contribute substantially to global primary production, yet many algal lineages still lack reference genomes with the continuity and annotation quality required for fine-scale structural, regulatory and comparative analyses. *Ostreococcus tauri*, one of the smallest known free-living photosynthetic eukaryotes, has been a model marine picoeukaryote for over two decades. Despite successive improvements to its historical reference genome, previous assemblies retained hundreds of gaps and incomplete genes, hampering high-resolution genomic analyses. Here, we present *O. tauri* RCC4221 genome version 2026, a telomere-to-telomere assembly of all 20 chromosomes spanning 13.34 Mb with no gaps. This assembly combines PacBio long-read sequencing, Illumina short-read polishing, correction of unresolved regions guided by independent Nanopore-based assemblies. The updated reference supports a curated annotation comprising 7,683 protein-coding genes, 48 tRNA genes, 3 rRNA operons, 116 medium-sized noncoding RNAs, one signal recognition particle RNA and 138 small nucleolar RNAs. It also improves gene-model integrity and recovers candidate coding loci absent from the 2014 reference. Structural analyses resolved the organization of the two atypical low-GC chromosome 2 and 19 that contain duplicated regions that were collapsed or misrepresented in previous assemblies. Finally, bisulfite sequencing and PacBio SMRT sequencing revealed a dual DNA methylation landscape, with CG-context cytosine methylation concentrated in gene bodies and N6-methyladenosine (m6A) enriched at the start codon. The updated *O. tauri* 2026 assembly provides a complete and curated reference resource for chromosome biology, comparative genomics, epigenomics and RNA biology in a model marine picoeukaryote.

## 1. Introduction

Marine phytoplankton are central to ocean biogeochemistry and account for a large fraction of primary production [1]. Genome sequences have become fundamental resources for connecting algal diversity with functional potential, phylogenomics and comparative biology [2]. However, algal genome quality remains uneven, and many assemblies still contain gaps, fragmented scaffolds or incomplete annotations that limit comparative and functional interpretation [3]. For compact unicellular algae, these limitations are especially consequential because coding genes, regulatory regions and noncoding RNAs are densely packed within short genomic intervals.

*Ostreococcus tauri* is among the smallest known free-living photosynthetic eukaryotes, with a cell diameter of approximately 0.8 μm, and one of the few experimental models among marine green picoalgae [4–7]. Its small cell size, compact genome, and ease of cultivation have made it a valuable system for studying the organization of minimal eukaryotic cells, photosynthetic acclimation [8], and host–virus interactions [9]. In addition, its phylogenetic position within the Mamiellophyceae makes it informative for comparative analyses of biosynthetic pathways, including taurine biosynthesis [10] and phytohormone synthesis [11], as well as chromosome evolution at the base of green lineage [7].

The first genome assembly of *O. tauri* was published in 2006 and represented a landmark achievement as the first sequenced genome of a marine green picoalga [5]. That assembly revealed several intriguing genomic features, including high gene density, compact intergenic regions, and the presence of two compositionally atypical chromosomes [5]. A later Illumina-based improvement of this reference assembly substantially reduced the number of gaps and improved annotation quality, but 477 unresolved regions, encoded as 20 consecutive ‘N’ in the chromosome sequences remained [12]. Repetitive DNA, duplicated segments, and low-complexity regions are particularly difficult to reconstruct accurately, affecting not only assembly completeness, but also the biological interpretation of chromosome architecture, gene content, and regulatory landscapes [13–16].

In *O. tauri*, the need for an improved reference genome is especially relevant because several of its most biologically interesting features are concentrated in structurally atypical genomic regions. Chromosome 19, the small outlier chromosome (SOC), is compositionally distinct from the 18 standard chromosomes and has been linked to virus-related phenotypic variation and extensive transcriptional [17] and structural plasticity in *O. tauri* [6]. Chromosome 2, the big outlier chromosome (BOC), contains a low-GC outlier region that is conserved across Mamiellales species with whole-genome sequences [18,19] and includes the candidate mating-type locus in this lineage [7].

The same reference-quality problem extends to annotation and regulation. Medium-sized noncoding RNAs and small nucleolar RNAs (snoRNAs) have already been shown to form a substantial component of the *O. tauri* transcriptome [20]. Yet noncoding RNA (ncRNA) discovery, coding-gene completeness and methylation profiling all depend on accurate coordinates around genes, repeats and duplicated segments. Cytosine methylation (5mC) in small marine eukaryotes can exhibit nucleosome-scale periodicity [21], whereas N6-methyladenosine (6mA) has been reported near the transcription start sites of actively transcribed genes in *Chlamydomonas* [22]. A fully resolved *O. tauri* genome therefore creates the opportunity to connect assembly correctness with gene annotation, ncRNA biology and epigenomic organization in the same reference framework.

Here, we present *O. tauri* 2026, a gapless telomere-to-telomere chromosome-scale reference for *O. tauri* RCC4221. We use this assembly to address three connected questions. First, which previously unresolved or collapsed regions prevented the historical reference from representing all chromosomes accurately? Second, how does a complete coordinate system improve the interpretation of protein-coding and ncRNA annotation? Third, what is the genome-wide cytosine and adenine methylation patterns on a gapless assembly? By linking assembly improvement to structural, annotation and methylation outcomes, this study positions *O. tauri* 2026 as a reference resource for future work on marine picoalgal genome biology.

## 2. Methods

### 2.1 Genomic Data

DNA extracted from strain RCC4221 was sequenced by Eurofins using the PacBio RSII technology with an 8-12 kb insert library. A total of 1.44 Gbp of PacBio reads were assembled with HGAP3 [23], producing a 13.42 Mbp draft assembly comprising 19 telomere-to-telomere (T2T) chromosomes. The only chromosome not assembled T2T was chromosome 16; therefore, the 528 Kbp sequence of chromosome 16 from the 2014 reference assembly [12], which contains 17 internal gaps, was retained as the starting sequence for this chromosome.

This initial assembly was polished with Pilon [24] using available DNA-seq data mapped to the assembly (Supplementary Table S1), applying base-level correction only (--fix bases), with a minimum read depth (mindepth) of 20 and a minimum quality (minmq) of 30. Following polishing, the resulting starting assembly length remained at 13.42 Mbp, comprising 19 T2T chromosomes.

Four assemblies of *O. tauri* RCC4221 were used for comparison and improvement of this starting assembly. The first two are the historical 2006 assembly (*O. tauri*_2006: 12.56 Mb, 1671 gaps, GenBank accession number NC_014426.1 to NC_014445.1), and the 2014 Illumina-improved assembly (12.92 Mbp, 477 gaps, GenBank accession number NC_014426.2 to NC_014445.2). These two previous assemblies are not independent as the 2014 version was generated by improving the 2006 version. Two additional and independent Nanopore assemblies of the same strain were also used, one adding up to 13.29 Mbp and 27 contigs sequenced at the CNAG (Barcelona) as previously described [25]. The second was sequenced at Bio2mar (Banyuls sur mer), and consisted of 19 contigs totaling 12.98 Mbp. Briefly, raw Nanopore reads were quality-checked with NanoPlot v1.44.1 [26] before and after adapter trimming with Porechop v0.2.4 [27]. Cleaned reads were assembled using Canu v2.3 [28]. Genome assembly was performed using the trimmed Nanopore reads with the following parameters: minOverlapLength= 1000 (minimum overlap required between reads), utgErrorRate=0.01 (error rate threshold for unitig construction), correctedErrorRate= 0.095 (expected error rate after read correction), and genomeSize= 14.3m (estimated genome size). Canu was executed in non-grid mode using 32 threads.

The number of non-canonical bases (i.e., non-ATGC bases) per sequence was estimated using the (*which*(!(*seq* %in% c(“a”, “t”, “g”, “c”))) function from the *seqinR* package [29].

### 2.2 Genome Polishing and Gap Resolution

The starting Pilon polished PacBio version contained 11,529 non-ATGC nucleotides. To resolve these sites, previously generated Illumina paired-end reads from *O. tauri* RCC4221 (SRA accessions: SRX026855 and SRX030853) were aligned to the assembly using BWA-MEM with default parameters [30]. Non-ATGC sites were replaced with the most frequently observed nucleotide, provided that at least 3 reads with the same base covered the site.

The remaining non-ATGC nucleotides showed a non-uniform distribution across the assembly, with many occurring in clusters. To resolve these sites systematically, we defined unresolved regions (UR) as regions containing adjacent ambiguous positions separated by less than 100 nucleotides. Adjacent URs were kept as separate regions only when separated by more than 500 canonical nucleotides (A, C, G, or T); otherwise, they were merged into a larger UR. For each UR, 500-bp flanking sequences upstream and downstream were extracted and aligned against two independent genome assemblies (Supplementary table S1) using BLASTn [31]. A UR was considered resolvable when: (i) the upstream and downstream flanking sequences both aligned to the same chromosome; (ii) each flanking sequence produced an alignment longer than 495 bp with sequence identity greater than 98%; and (iii) the two alignments were in the same orientation. If these conditions were satisfied, the corresponding sequence spanning the upstream and downstream alignments was extracted and used to replace the UR in the assembly. When the different independent genome assemblies provided different lengths for UR, we arbitrarily chose the solution closest in length to the corresponding UR.

For the remaining URs, including those on chromosome 16, we used a dotplot visualization approach using NUCmer from the MUMmer4 package [32]. Entire chromosomes containing URs were aligned against the corresponding reference genome chromosomes to generate delta alignment files. These delta files were subsequently visualized with Assemblytics [33]. This analysis revealed assembly errors in the previous version of chromosome 16, including two translocations and four segmental duplications, which accounted for the URs flanking these misassembled regions.

To test the recent SOC chromosome evolution model hypothesis inferred in the sister species *O. mediterraneus* [25], the updated version of SOC chromosome 19 version was screened for signatures of translocations from other chromosomes. Non-overlapping 500-bp windows were generated from the SOC and aligned against the standard chromosomes, including chromosome 2, using BLASTn with the parameters -dust no -soft_masking false to keep low complexity regions included. Only unique hits between chromosome 19 and a single other chromosome, with >90% sequence identity over at least 500 bp, were retained. This filtering strategy was used to exclude transposable and repetitive elements, which are abundant on chromosomes 19 and 2, and also occur on other chromosomes in order to highlight putative signatures of past chromosomal fusions rather than transposable element activity. The retained hits were visualized in R using the circlize package [34].

### 2.3 Genome Annotation update

To annotate the updated genome, we first transferred annotations from the 2014 reference genome to the updated assembly using Liftoff [47] with default parameters. Following the transfer, we identified 426 problematic annotations with missing start codons, missing stop codons or inframe stop codons. To correct these gene predictions, RNA-Seq reads (Supplementary Table S1) were aligned to the updated genome using HISAT2 [48] with the parameter --max-intronlen 5000. Gene annotations and RNA-Seq alignments were visualized and inspected using GenomeView [49], allowing simultaneous examination of gene models, supporting transcript evidence, and manual gene-structure corrections. Second, genomic regions present in the updated assembly but absent from the 2014 reference genome were identified by whole-genome alignment using minimap2. These additional regions were extracted using BEDTools and searched for potential protein-coding genes using BLASTX against the NCBI non-redundant protein database. Candidate genes were further evaluated using RNA-Seq read alignments and InterProScan domain annotation.

Third, transfer RNA (tRNA) genes were identified using tRNAscan-SE v2.0 [50] with default eukaryotic parameters. We also searched for permuted tRNAs, which have been reported in *O. tauri* [51], using custom BLAST-based approaches with known permuted tRNA sequences as queries.

Fourth, ribosomal RNA (rRNA) genes were annotated using RNAmmer v1.2 [52] configured for eukaryotic rRNA prediction. Because RNAmmer does not detect 5.8S rRNA, 5.8S sequences were additionally identified using Infernal v1.1.4 [53] with covariance models from the Rfam database [54].

The complete annotation set was processed and validated using the AGAT toolkit (Another Gtf/Gff Analysis Toolkit) [55]. This step included identifying format inconsistencies, resolving structural errors, and generating summary statistics to ensure annotation accuracy and standard compliance. The finalized GFF3 file was subsequently converted into EMBL format using emblmygff3 [56].

### 2.4 RNA Sequencing for medium-sized ncRNA annotation

Cultures of *O. tauri* RCC4221 in exponential growth phase (cell density 2×10^7^ cells/ml) were sampled at 10h30 and 14h30 post-dawn, with three biological replicates per time point and 200 mL of culture per replicate. These time points were selected to cover *O. tauri’s* circadian cycle and maximize RNA yield and quality. Samples were centrifuged at 8,000 × g for 20 min. Cell pellets were resuspended in Tri-reagent buffer from the Direct-Zol kit, subjected to mechanical lysis (TissueLyser II, Qiagen, 2×45s at 30 Hz), and RNA was extracted following the manufacturer’s protocol. RNA concentration was measured by fluorometry (Quantus, Promega) and RNA integrity evaluated with Agilent 2100 Bioanalyzer. Library preparation included size selection to retain 50–300 nt fragments and enrich for small RNAs and Cap-Clip pyrophosphatase treatment to remove 5′-m7G caps and enable sequencing of previously capped RNAs. Illumina NextSeq paired-end sequencing (2 × 75 bp) was performed by Fasteris (Supplementary Table S1).

Samples from both time points were pooled by biological replicate, generating three independent biological replicate libraries for downstream analyses. Sequencing quality was assessed with FastQC (v0.11.9) [35], and low-quality sequences and Illumina adapters removed with Trim Galore (v0.4.5) [36]. Quality-filtered reads were mapped to the genome using STAR v2.6.0c [37] with key parameters: --outMultimapperOrder Random (random ordering of multi-mappers), --outFilterMultimapNmax 12 (maximum 12 alignment positions), --outFilterMismatchNoverLmax 0.02 (≤2% mismatch rate), --alignIntronMin 10 --alignIntronMax 500 (intron length range 10-500 bp). Because repetitive regions in the *O. tauri* genome produced multiple alignments for some reads, Samtools [38] was used to retain primary alignments and remove secondary or supplementary alignments.

Alignments were first visually inspected with IGV [39] to confirm expression of previously identified ncRNA loci. Alignments were then assembled into candidate transcripts using StringTie [40], and GffCompare [41] was used to identify transcripts absent from the reference annotation. Candidate transcripts were manually inspected (Supplementary Fig. S1) and filtered according to criteria established by Bousquet et al. [20], including stable expression across multiple samples, clear transcription start and end boundaries, and absence of obvious sequencing or alignment artifacts.

Functional Annotation. Retained candidate sequences were evaluated for coding potential using CPC [42], with CPC scores <0 considered indicative of RNA-coding transcripts. snoScan [43], and snoReport [44] were used to predict structural features of tRNAs and snoRNAs. Additionally, BLAST searches (E-value < 10^-5^) against the snoRNA ortholog database SnOPy [45] and miRbase v22 [46] were used to identify potential homologs of known small RNA families and conserved hairpin structures.

Representative candidate RNAs were validated by Northern blot. Total RNA (2-6 µg) was separated on 8% polyacrylamide gels (7×11 cm), with DNA oligonucleotides (25, 50, 75, 100 nt) and U1 RNA (163 nt) as size markers. RNA was electrotransferred to Hybond-NX membranes and hybridized overnight with sequence-specific oligonucleotide probes labeled with γ-32P-ATP using T4 polynucleotide kinase in ULTRAhyb-Oligo buffer. Membranes were washed at 50°C with low-stringency buffer (2 × SSC, 0.1% SDS), followed by high-stringency buffer (0.1 × SSC, 0.1% SDS), for 15 min each. X-ray films were exposed for 3–15 h, with or without intensifying screens depending on signal intensity.

### 2.5 Bisulfite and PacBio methylation analyses

Cultures of *O. tauri* strain RCC4221 were maintained in 30 ml L1 medium at 20°C under 12h light/12h dark cycles [17]. Cells were processed 5 hours after the onset of the light period when reaching 10^7^ cells/ml. DNA extraction was performed as described in Blanc-Mathieu et al. [12]. Bisulfite sequencing libraries were prepared using the Bioscientific NextFlex kit and sequenced at the GeT-Plage platform in Toulouse with Illumina HiSeq3000 system (paired-end 2×150 bp). Sequencing reads were mapped to the reference genome using Bismark [57]. Positions covered by fewer than 10 reads were excluded from downstream analyses.

Four *O. tauri* RCC4221 cultures were subjected to PacBio single-molecule real-time long-read sequencing [6]. PacBio sequencing can detect DNA modifications, including N6-methyladenine, through changes in DNA polymerase kinetics, particularly interpulse duration (IPD) and IPD ratio, following the principle described by Flusberg et al. [58].

All data processing and visualization were performed in R [59] with the ggplot2 package [60].

## Results

### 3.1 The integration of different methods was necessary for achieving completeness of the genome assembly

The starting PacBio-based assembly represented a major improvement in continuity over the previous *O. tauri* 12.92 Mbp reference [12], including telomere to telomere reconstruction of the atypical chromosomes 2 and 19. However, continuity alone was not sufficient to produce a complete reference. The starting assembly contained 11,529 non-ATGC positions, thereafter “ambiguous” positions, across a total assembly length of 13.42 Mb. Illumina-supported correction resolved 1,481 (13%) of these positions (Supplementary Table S2). The remaining 10,048 ambiguous positions could be clustered into 178 unresolved regions (URs) ranging from 1 to 3,387 bp and each containing at least one non-ATGC nucleotide (see Methods). By using flanking sequences of the URs to locate the corresponding intervals in independent *O. tauri* assemblies, we successfully resolved 170 URs (Supplementary Table S3). The eight remaining URs required further investigation.

Two URs corresponded to single ambiguous ‘N’ positions on chromosome 2, where DNA-seq coverage was unexpectedly low or absent. Although the cause of this local lack of DNA-seq support remains unclear, both positions were resolved during annotation refinement with support from RNA-seq evidence. The remaining six URs were located on chromosome 16. Comparison with an independent Nanopore assembly (A1662, Accession number: S-BSST3163) showed that the UR’s flanking regions mapped to different chromosomes, indicating that these regions reflected structural errors in the reference version of chromosome 16 rather than simple missing bases (Supplementary Fig. S2). Correction of chromosome 16, including deletion of duplicated regions and removal of two spurious insertions derived from chromosome 1, resolved all six URs. In addition, one missing telomeric tract on chromosome 16 was also recovered by alignment to an independent Nanopore assembly (Canu, Accession number: S-BSST3163) and incorporated into the final chromosome version. These steps resolved all 178 URs and eliminated the remaining non-ATGC positions from the final assembly.

### 3.2 Main improvements of the updated genome assembly

These corrections produced the updated reference genome assembly *O. tauri* 2026, a 13.34 Mbp reference composed of 20 gapless chromosomes, representing a 420 kb increase relative to the previous reference while eliminating all assembly gaps (Table 1). Consistent with this genome-level completeness, BUSCO analysis using the chlorophyta_odb10 dataset recovered 98.6% of the 1,519 expected chlorophyte orthologs as complete BUSCOs.

**Table 1.** Genome features of the three available genome versions of *O. Tauri* RCC4221.

|  | 2006 | 2014 | 2026 |
| --- | --- | --- | --- |
| genome length (Mbp) | 12.56 | 12.92 | 13.34 |
| Gaps | 1,671 | 477 | 0 |
| CDS | 8,166 | 7,661 | 7,683 |
| Complete CDS | 6,265 (76.7%) | 7,462 (97%) | 7,683 (100%) |
| tRNAs | 39 | 39 | 48 |
| rRNA | 1 | 1 | 3 |
| T2T chromosomes | 10 | 10 | 20 |
| 1T chromosomes | 10 | 10 | 0 |
| mncRNAs | 0 | 0 | 116 |
| SRP RNAs | 0 | 0 | 1 |
| snoRNAs | 0 | 0 | 138 |

Whole-genome dotplot analysis between the 2026 and the 2014 Illumina-based assembly confirmed high global collinearity, with all chromosomes exhibiting clear one-to-one correspondence (Fig. 1A). The main differences were concentrated in chromosomes 2 and 19, consistent with previous reports that these chromosomes are compositionally atypical and structurally unusual [5]. In particular, large duplications on chromosomes 2 and 19 that were collapsed in the 2014 assembly are now resolved. The lower GC region is now located centrally on the updated chromosome 2 (Fig. 1B), consistent with the structure of this chromosome in all other Big Outlier Chromosomes (BOCs) of Mamiellales including 2 *Micromonas*, 3 *Ostreococcus*, and 1 *Bathycoccus* [7]. The new assembly therefore provides a more accurate sequence framework for future comparative analyses of low-GC, sex-related outlier chromosomes in Mamiellophyceae.

**Fig. 1.**
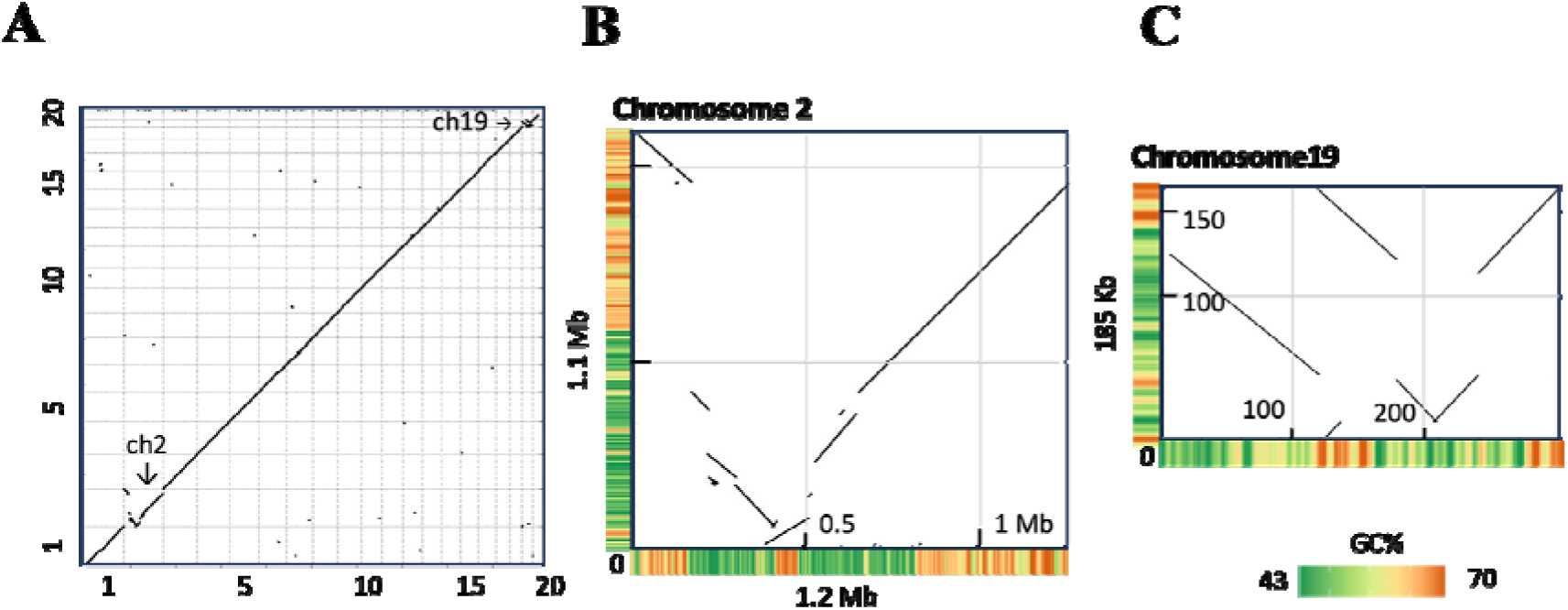
Whole-genome and chromosome-specific assembly analysis between the previous *O. tauri* 2014 and the updated *O. tauri* 2026 versions. **A.** Genome-wide dotplot showing overall synteny; **B.** Detailed synteny of chromosome 02 (BOC); **C.** Detailed synteny of chromosome 19 (SOC).

Chromosome 19,the small outlier chromosome (SOC), was reassembled by resolving an internal duplication that had been collapsed into a single region in the 2014 assembly (Fig. 1C). Screening this updated chromosome 19 for signatures of past translocations from other chromosomes (excluding transposable elements, see method section 2.2), we identified 11 regions larger than 500 bp shared with standard chromosomes 9, 11, 16, 17 and 20, as well as 18 blocks shared between chromosomes 19 and 2 (Fig. 2). Several chromosome 19 intervals shared with standard chromosomes overlapped annotated coding sequences, including transporter-domain genes, methyltransferase-family genes, leucine-rich-repeat or surface-repeat proteins, and CDSs annotated with reverse-transcriptase/RNA-dependent DNA polymerase domains.

**Fig. 2.**
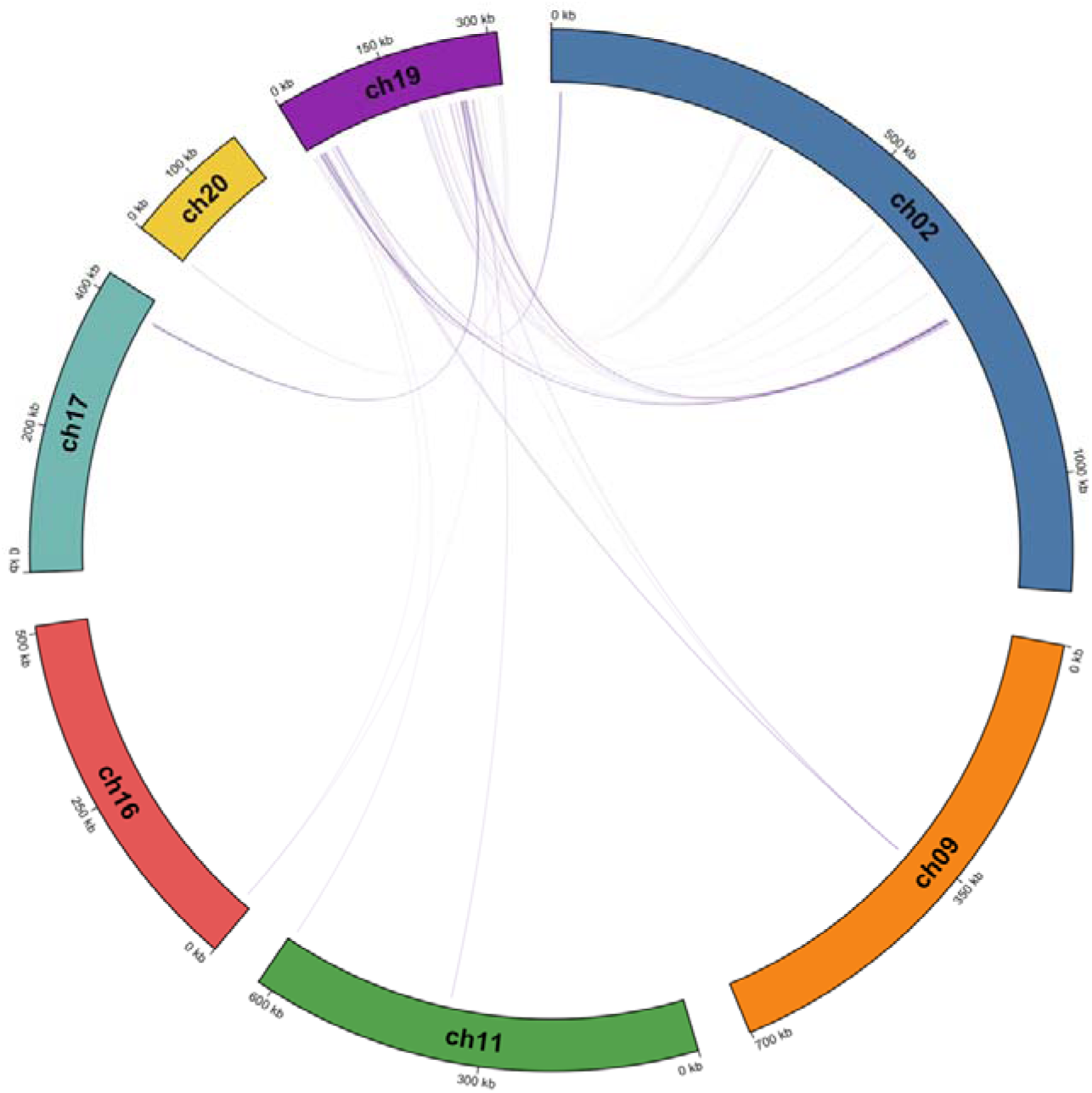
Circos plot of homologous regions shared between chromosome 19 (SOC) and other chromosomes in *O. tauri*.

Across the updated assembly, improved genome completeness also recovered eight candidate protein-coding loci longer than 500 bp that were absent from the 2014 reference (Supplementary Table S4). These loci were supported by RNA-seq evidence, protein similarity and conserved domains, indicating that the updated assembly restored not only repetitive or structurally complex regions but also previously missing coding sequence. Their predicted annotations included RNA- or ribosome-associated proteins, as well as candidates with metabolic or surface-associated functions.

### 3.3 Genome Annotation and Curation

Annotation transfer using Liftoff successfully mapped 7,587 genes from the *O. tauri* 2014 annotation to the updated genome, whereas 74 gene models could not be transferred. Manual inspection of these 74 unmapped gene models revealed multiple causes for the transfer failures, including incomplete or erroneous gene models in the previous annotation, putative redundant copies, and loci requiring RNA-seq-guided correction (Supplementary Table S5). Quality assessment of the transferred annotations identified 426 genes with structural errors, such as in-frame stop codons, missing start or stop codons, and incomplete gene boundaries; these were manually corrected in GenomeView [50] using RNA-Seq evidence. The corrections restored gene continuity in regions where unresolved sequence or local assembly errors had previously interrupted coding models.

To further evaluate the improvement achieved by this annotation update, we compared CDS integrity between the previous and updated annotations. The 2014 annotation contained 298 unique affected protein-coding models, including 148 incomplete models lacking a valid start and/or stop codon, 123 models containing internal stop codons, and 27 models affected only by ambiguous or gap-containing CDS sequence. By contrast, all 7,683 protein-coding sequences in the updated annotation passed these basic CDS-integrity checks, indicating that the updated reference and manual curation improved gene-model continuity as well as genome contiguity.

To complement our understanding of the *O. tauri* transcriptional and regulatory landscape, we integrated the previously reported large family of snoRNA candidates (138), into the complete genome coordinate system [20]. The updated annotation also contains 116 medium-sized noncoding RNAs, one SRP RNA, 48 tRNA genes and three rRNA operons (Table 1). The inclusion of these ncRNA families substantially enriches the functional landscape of the *O. tauri* genome, providing a valuable resource for investigating RNA-based regulatory mechanisms in this model picoalga. Together, these annotations provide an integrated genomic resource that links ncRNA loci to intronic, intergenic, dicistronic tRNA-associated and antisense contexts within a complete chromosome-scale coordinate system (Table 2). The validation of two predicted medium-sized ncRNAs by Northern blot using specific radioactive probes on RNA extracted at two different times within the 24h cycle is presented Fig. 3A. First, mncR37, which is present in two copies in the genome (Fig. 3B and D). Second, the dicistronic mncR67-tRNA^Glu^ (Fig. 3C and E), for which mncR67 (Fig. 3E), as well as the precursor mncRNA containing both the tRNA and mncR67 (Supplementary Fig. S3) could be validated. Because RNA-seq libraries were prepared in two conditions (Fig. 3A), this hybridization approach could even detect a difference in expression of Ot-mncR37 and Ot-mncR67 between control and virus-infected culture conditions, namely a lower expression of both mncRNAs in the infected cultures (Fig. 3D and Fig. 3E).

**Fig. 3.**
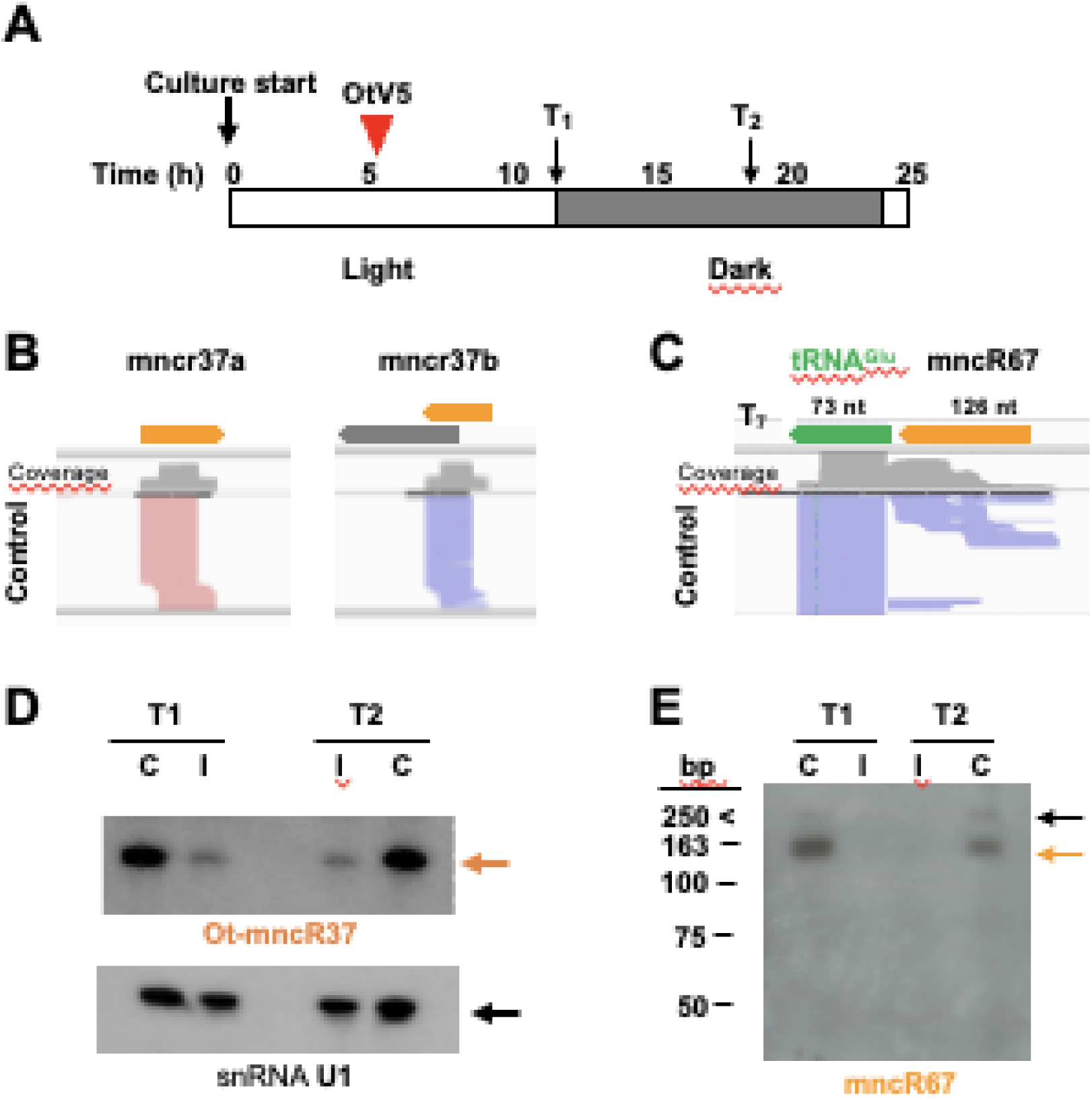
Validation of predicted mncRNA by Northern blot using specific probes. **A.** Experimental design to produce the targeted mncRNA libraries. Clonal cell cultures were grown under a 12h light/12h dark regime under two conditions; control (C) and infected (I) by prasinovirus OtV5. Time (h) indicates hours since the beginning of the light period. T_1_ and T_2_, indicate sampling 4h30 and 8h30 post infection by prasinovirus OtV5, for which RNA sequencing was performed in I and C. **B.** Read profiles of Ot-mncR37a and Ot-mncR37b isoforms. Ot-mncR37b overlaps with the 5’end of gene OTAU_ost16g00010 (gray arrow) predicted to encode a small protein of unknown function. **C.** Read profiles of mncR67-tRNA^Glu^ gene. **D.** Northern blot analysis of Ot-mncRNA37 expression (Probe sequence: TCGACGCTGTCGATGACTTCTACC, orange arrow). U1 snRNA was used as a control, detected with a specific probe (black arrow). **E.** Northern analysis of mncR67-tRNA^Glu^ gene expression. The black arrow indicates the dicistronic precursor. The Ot-mncR67 signal at the expected size is indicated by the orange arrow (Probe sequence: CGATGTATTCCGTGCGCTAGTAGAG).

**Table 2.** Genomic context of mncRNA loci in.

| Genomic Organization | Ot-mncRNAs | Ot-SRP | Ot-snoRNAs |
| --- | --- | --- | --- |
| Intronic | 61 | 0 | 84 |
| Intergenic | 42 | 1 | 15 |
| Clustered | 2 | 0 | 37 |
| Dicistronic tRNA | 10 | 0 | 0 |
| Antisense to CDS | 1 | 0 | 2 |
| Total | 116 | 1 | 138 |

### 3.4 The cytosine methylation (5mC) landscape in *O. tauri*

To characterize the epigenetic landscape of *O. tauri*, we performed bisulfite sequencing and PacBio SMRT sequencing to detect cytosine and adenine methylation across the genome. Overall, 11.8% of cytosines (±0.16%, n=11) were methylated as 5-methylcytosine (5mC) in the *O. tauri* genome. Notably, cytosine methylation was almost exclusively restricted to the CG dinucleotide context, with 99.995% (±0.0015%, *n*=11) of 5mC occurring as 5mCG, indicating minimal CHG or CHH methylation typical of plant genomes.

Examining 5mCG methylation across different genomic contexts revealed functional compartmentalization. Intergenic regions displayed 12% methylation of CG sites, while protein-coding sequences (CDS) showed significantly higher methylation at 27% of CG dinucleotides. Within coding sequences, methylation exhibited a characteristic positional bias: methylation rates were markedly depleted around transcription start sites (TSS), reaching a minimum near the ATG start codon (∼6% methylation rate), then rapidly increased within the first 200-300 bp of the coding sequence, plateauing at approximately 31-33% throughout the gene body before declining slightly toward the transcription termination site (Fig. 4). This gene body methylation pattern, with pronounced TSS depletion, suggests a role for 5mCG in regulating transcriptional elongation or co-transcriptional processes rather than transcriptional initiation in this compact genome.

**Fig. 4:**
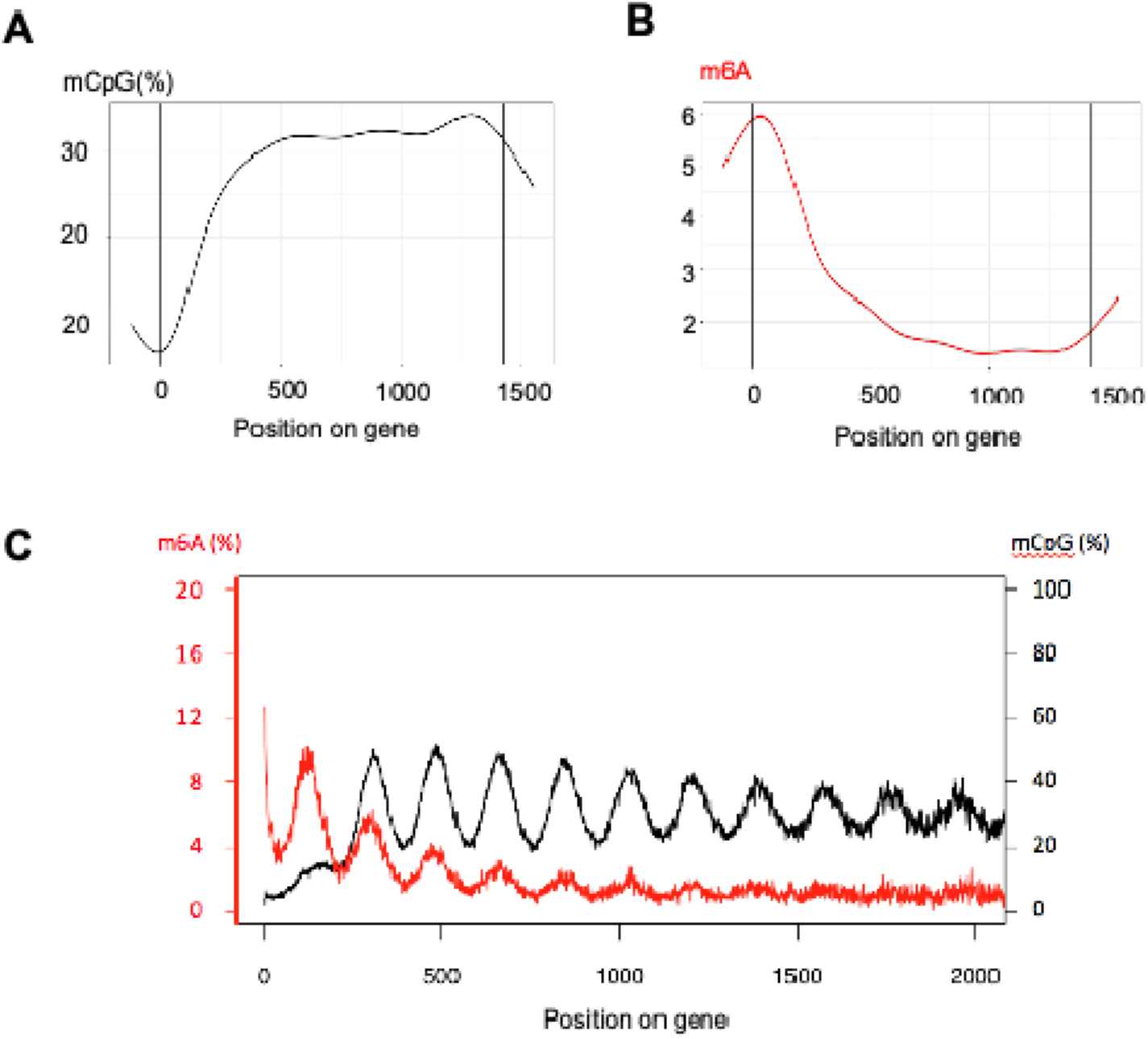
Gene body methylation in *O. tauri*. Average 5mCG methylation rate in and around protein coding genes. Positions of cytosines in genes were calculated by their relative distances to the start codon. Genes lengths have been adjusted proportionately to the average length of all genes. B. Average 6mA methylation rate in and around protein coding genes. Positions of adenine in genes were calculated by their relative distances to the start codon. Genes lengths have been adjusted proportionately to the average length of all genes. C. Average 6mA and 5mCG methylation in absolute distance from start ATG (position 0).

### 3.5 The adenine methylation (6mA) landscape in *O. tauri*

In contrast to cytosine methylation, adenine methylation (6mA) was relatively sparse across the genome, with only 2.8% of adenines modified. However, 6mA distribution was non-random and displayed a pattern inversely correlated with 5mCG (Fig. 4). Strikingly, 6mA was enriched at transcription start sites, with the highest peak occurring at the ATG start codon itself. Specifically, adenines near the start codon showed elevated methylation frequencies: the first adenine near ATG was methylated at 13.75%, and the second adenine at 13.26%. This 6mA enrichment rapidly declined within the first 100-200 bp of the coding sequence, reaching baseline levels (∼1.5-2%) throughout the gene body before showing a modest increase near the 3’ end of genes.

The complementary distributions of 5mCG (gene body-enriched, TSS-depleted) and 6mA (TSS-enriched, gene body -depleted) suggest these two methylation marks may play distinct and potentially antagonistic roles in transcriptional regulation in *O. tauri*.

The data revealed a striking transition zone at gene starts: 6mA peaks sharply in the start ATG codon region, while 5mCG shows reciprocal dynamics, with minimum levels at the TSS and maximum levels in the gene body (mCpG ∼30-40%) (Fig. 4). This inverse relationship creates distinct epigenetic domains that may demarcate regulatory boundaries between promoters and transcribed regions.

## 4. Discussion

### 4.1 From contiguity to correctness: refining genome assemblies in the long-read era

Genome assemblies have historically relied on short-read sequencing technologies such as Illumina, which provide high base-level accuracy and cost efficiency and have therefore been widely used for genome polishing and variant detection [61]. However, the short read length restricts their ability to resolve repetitive and structurally complex regions, often resulting in assembly gaps, repeat collapse, and mis-joins, particularly in repeat-rich or compositionally biased genomic regions [15,16]. Recent advances in long-read sequencing have substantially improved both read length and accuracy, with reported base-level accuracies exceeding 99.5–99.9% [62,63], enabling near-complete and T2T genome assemblies [64].

The benefits of long-read resequencing were most evident on chromosomes 2, 16 and 19. Chromosomes 2 and 19 contain compositionally atypical and structurally complex regions that were difficult to represent accurately in the previous short-read-improved reference. On chromosome 2, the updated assembly corrected the placement and structure of features within the low-GC region. On chromosome 19, it resolved duplicated segments that had previously been represented as single-copy, consistent with repeat collapse or local misrepresentation in the 2014 assembly. The updated chromosome 16 sequence further corrected errors in the previous assembly by identifying and removing spurious insertions of sequences from chromosome 1. Together, these examples show that the main improvement of the updated reference is not only gap removal, but also the correction of assembly errors.

Despite this, long-read sequencing alone does not automatically guarantee a fully resolved reference genome. Even highly contiguous assemblies can retain local errors or weakly supported regions in duplications, low-complexity sequences and repeat-rich intervals, making independent assessments of assembly correctness essential [16,65]. This is consistent with our observation that a substantial number of ambiguous non-ATGC positions persisted in the initial assembly despite high overall continuity. Resolving these regions required the integration of multiple independent data sources, including short-read validation [66], transcriptomic evidence [16], comparisons with several independent long read assemblies [65].

Taken together, our results emphasize that genome assembly quality in the long-read era is no longer primarily limited by contiguity, but by the accurate resolution and validation of complex genomic regions [16,64]. In *O. tauri*, this distinction has direct consequences for genome annotation, as regions absent or poorly represented in the 2014 reference affected both gene-model integrity and the recovery of coding loci [12]. By resolving these regions in a complete chromosome-scale coordinate system, the updated assembly provides a more reliable foundation for comparative genomics, chromosome biology and functional annotation in this model picoalga.

### 4.2 Structural characterization of chromosome 19 in the updated assembly and its implications for virus-related adaptation

The updated PacBio assembly refines the structural description of chromosome 19, the small outlier chromosome (SOC) of *O. tauri*. This chromosome has been recognized since the first genome publication as compositionally distinct from the standard chromosomes and was later associated with marked intraspecific variability and virus-related phenotypes [5,6,12]. Compared with the 2014 Illumina-based reference, regions previously represented as single-copy are now resolved as duplicated segments, indicating that parts of the SOC were collapsed in the earlier assembly. In addition, a NUCmer-based self-alignment of the SOC revealed at least four nonredundant internal inverted duplicated modules larger than 5 kb, represented by paired intervals of 11.9, 7.4, 45.8, and 30.4 kb (Table 3). These observations indicate that the structural complexity of the SOC is expressed not only as chromosome-scale size heterogeneity, as described in natural populations [6], but also as internal architectural complexity involving duplicated and inversion-related modules.

**Table 3.** Internal inverted duplicated modules identified on ch19.

| Block ID | Representative interval on ch19 | Paired interval on ch19 | Length (kb) | Identity (%) |
| --- | --- | --- | --- | --- |
| B1 | 6567–18497 | 251710–263649 | 11.9 | 99.46 |
| B2 | 123120–130557 | 304681–312118 | 7.4 | 99.77 |
| B3 | 138784–184596 | 260239–306039 | 45.8 | 99.88 |
| B4 | 184882–215245 | 215661–246026 | 30.4 | 99.98 |

Annotation of the four major inverted blocks revealed that these regions carry multiple coding sequences with non-random functional content. The two largest blocks (B3, 45.8 kb; B4, 30.4 kb) were enriched in enzymes associated with carbohydrate modification and cell-surface biosynthesis, including galactosyltransferases, rhamnan synthesis-related proteins, NAD-dependent epimerase/dehydratase, and CMP-N-acetylneuraminic acid hydroxylase. Earlier work on virus-resistant *O. tauri* lines reported differential expression of multiple glycosyltransferase genes on this chromosome, prompting the hypothesis that altered carbohydrate or surface modification may affect host–virus interactions, including viral adsorption or later stages of the infection cycle [17,67]. In addition, several blocks also contained annotations associated with mobile-element or rearrangement-related functions, including a reverse transcriptase, a Harbinger transposase-derived protein, and a DDE superfamily endonuclease, consistent with the established view that the SOC intraspecific hypervariability is generated by a high rate of duplications, translocations, and internal rearrangements [6,25].

### 4.3 DNA methylation landscape of the *O. tauri* genome

Cytosine methylation in *O. tauri* is almost exclusively restricted to the CG dinucleotide context (99.995% as 5mCG), placing this species closer to the predominantly CG-based methylation regime seen in animals than to the multi-context methylation landscape typical of land plants, which display prominent CHG and CHH methylation [69,70]. The characteristic depletion of 5mCG at transcription start sites (TSS, minimum ∼6% at the ATG codon), followed by rapid enrichment across the coding region (plateauing at ∼31–33%), is broadly consistent with patterns described in other eukaryotes and is often associated with stably or broadly expressed genes [71,72]). In the extremely compact *O. tauri* genome, where intergenic distances are among the shortest in free-living eukaryotes [5], such TSS-proximal hypomethylation may help maintain promoter regions permissive to transcription factor binding, whereas gene body methylation may contribute to the suppression of inappropriate transcription initiation within coding regions [73]. Notably, the 5mCG profile within gene bodies displays a periodic oscillation of approximately 180 bp (Fig. 4), corresponding closely to the nucleosome repeat length and consistent with preferential methylation of linker DNA over nucleosome-wrapped sequences, a pattern previously identified in *O. lucimarinus* and proposed as an ancient feature of chromatin organization in marine unicellular eukaryotes [21].

Adenine methylation (6mA), although relatively sparse overall (2.8% of adenines), shows a strikingly complementary distribution to that of 5mCG: it is sharply enriched at the TSS, with adenines near the ATG start codon methylated at 13.75% and 13.26%, respectively. Adenine methylation then declines rapidly to baseline levels within the first 100–200 bp of the coding sequence (Fig. 4). This TSS-enriched 6mA pattern parallels observations in *C. reinhardtii*, where 6mA marks active transcription start sites and has been proposed to influence nucleosome positioning at promoters [22]. The inverse relationship between 5mCG and 6mA along the gene axis, with 6mA enriched in the promoter-proximal region and 5mCG concentrated in the transcribed gene body, suggests that these two marks define distinct epigenetic domains rather than acting redundantly [74,75]. In one of the most gene-dense eukaryotic genomes known, the two distinct DNA methylation systems suggest a non-random partitioning of epigenetic information between promoter-associated and gene body-associated regions. We note, however, that the biological interpretation of 6mA in eukaryotes remains an active area of investigation, and the functional consequences of the patterns reported here will require further experimental investigation.

## 5. Conclusions

This complete and curated reference genome for *O. tauri* RCC4221 resolves gaps from previous assemblies and corrects structurally complex regions gene annotation. This assembly improves the resolution of the two atypical BOC and SOC chromosomes, coding-gene completeness, ncRNA organization and DNA methylation landscapes. It therefore establishes a strengthened reference for future comparative genomics, genome biology and epigenomics studies in marine picoeukaryotes.

## Credit authorship contribution statement

**Gaokang Liu:** Investigation, Genome assembly improvement, Annotation curation, Formal analysis, Visualization, Writing first draft, review & editing. **Laurie Bousquet:** RNA-seq preparation and analyses, mncRNA annotation, review & editing. **Helene Mayeur:** Bisulfite sequencing analyses, bioinformatic analyses, review & editing. **Eric Manirakiza:** Genome assembly, Data curation, review & editing. **Vladimir Daric:** Genome assembly, Data curation, review & editing. **David Lopez-Escardo:** Supervision, review & editing. **Céline Noirot:** PacBio adenine methylation analysis. **Christophe Klopp:** PacBio adenine methylation analysis, review & editing. **Sheree Yau:** Supervision, Study conception and design, Funding acquisition, review & editing. **Marc Krasovec:** Manual annotation curation, Data interpretation, review & editing. **Nigel Grimsley:** RNA-seq analyses, Funding acquisition, Supervision, review & editing**. Manuel Echeverria:** mncRNA study conception and design, mncRNA annotation, manual curation, review & editing. **Gwenaël Piganeau:** Study conception and design, Supervision, Funding acquisition, Genome annotation, Manual curation, Writing – original draft, review & editing.

## Acknowledgements

This paper is dedicated to Hervé Moreau, who passed away in July 2020. He made a major contribution to the field of phytoplanktonic marine models. We are grateful to all past and present members of the Genophy group for stimulating discussions and insightful comments, particularly Frédéric Sanchez, Anaïs Labecot, and Lisa Mettrop for their assistance with cultures and sequencing. We thank the GenoToul bioinformatics platform for access to computing facilities, and David Pecqueur and Christophe Salmeron at the BIOPIC platform for support with the cytometry facilities. We also thank Aurelie Claes and Nyree West from the BIO2MAR platform for Nanopore sequencing.

## Data availability

The genome assembly generated in this study has been deposited in the European Nucleotide Archive under BioProject accession PRJEB98424, with assembly accession GCA_977066015.1 and chromosome sequence accessions OZ477818.1–OZ477837.1. The two independent Nanopore assemblies used for validation, A1662 and Canu, are available in BioStudies under accession S-BSST3163, the complete list of accession numbers is available in Table S1.

## Declaration of generative AI use

During the preparation of this work, the author(s) used LeChat (Mistral AI) for english editing. The author(s) reviewed and edited the output as needed and take full responsibility for the content of the published article.

## Supplementary Tables

Supplementary Table S1. Datasets used for the updated reference genome

Supplementary Table S2. Illumina read-supported correction of 1481 non-ATGC positions

Supplementary Table S3. Unresolved regions solved by alignment of flanking sequences

Supplementary Table S4. Eight putative protein-coding loci longer than 500 bp from regions absent in the 2014 reference

Supplementary Table S5. Liftoff failed gene curation

## Supplementary Figures

Supplementary Fig. S1 Read mapping of the short read RNA-Seq libraries against the reference genome sequence for the predicted antisense noncoding RNA Ot-mncR9 (OTAU_ost02g01100).

Supplementary Fig. S2 Alignment of chromosome 16 against an independent assembly

Supplementary Fig. S3 Validation or expression of tRNA^Glu^ and response of dicistronic mncR67-tRNAGlu gene in two conditions and two timepoints: control (C) and post OtV5 infection (I).

